# Repetition effects in action planning reflect effector- but not hemisphere-specific coding

**DOI:** 10.1101/2021.07.16.450393

**Authors:** Christian Seegelke, Carolin Schonard, Tobias Heed

## Abstract

Action choices are influenced by future and recent past action states. For example, when performing two actions in succession, response times (RT) to initiate the second action are reduced when the same hand is used. These findings suggest the existence of effector-specific processing for action planning. However, given that each hand is primarily controlled by the contralateral hemisphere, the RT benefit might actually reflect effector-independent, hemisphere-specific rather than effector-specific repetition effects. Here, participants performed two consecutive movements, each with a hand or a foot, in one of two directions. Direction was specified in an egocentric reference frame (inward, outward) or in an allocentric reference frame (left, right). Successive actions were initiated faster when the same limb (e.g., left hand - left hand), but not when the other limb of the same body side (e.g., left foot - left hand) executed the second action. The same-limb advantage was evident even when the two movements involved different directions, whether specified egocentrically or allocentrically. Corroborating evidence from computational modeling lends support to the claim that repetition effects in action planning reflect persistent changes in baseline activity within neural populations that encode effector-specific action plans.

## Introduction

Interacting with our environment continually requires us to decide between multiple potential actions. Consider the simple decision of choosing an effector to open a door. Intuitively, one might choose between the left and right hand. However, when our hands are occupied, for example because we are carrying a stack of heavy folders, we can flexibly come up with another effector option and open the door with an elbow or knee.

Neurophysiologically inspired models such as the Affordance Competition Hypothesis regard the process of action selection as a competitive decision process between multiple, simultaneously prepared actions (1, 2). In this view, a distributed network of posterior parietal and premotor cortical regions continuously specifies all currently possible actions in parallel (3–6). Once the neural activity for one of the potential actions passes a threshold, the selected response is initiated.

Several factors can bias this decision process: In the context of choosing between different limbs (i.e. left vs. right hand), a decisional process subsumed under the domain of action planning (7), hand choice is influenced by handedness (8), future task demands (9), biomechanical costs (10, 11), spatial reference frames (12), and the recent motor history, also termed motor hysteresis (11, 13–15). Motor history effects manifest as a bias of hand choice for a new action towards the effector that was used for the previous action, especially when the biomechanical costs of responding for alternative effectors, such as the two hands, is similar. The prevailing explanation for the existence of motor hysteresis has been that it is computationally more efficient to reuse and slightly modify a previously used action plan than to create a new plan for each single action (16, 17).

Recent behavioral and neuroimaging data have provided support for this ‘planning efficiency account’ (18, 19). In these studies, participants performed two actions in succession, referred to as prime and probe actions, respectively. These actions involved grasping a tool with one hand and rotating it in one of two directions – either inward or outward with respect to the body midline – each with a distinct grasp. The study design comprised four conditions that reflected the relationship between prime and probe actions: Participants repeated the same action (identical repeat, IR); they used the same hand but executed different rotations and grasps (hand repeat, HR); they repeated the same rotation and grasp but used different hands in the two consecutive actions (grasp repeat, GR), or used both a different hand and rotation/grasp (no repeat, NR). Response time (RT) to initiate the probe action was shorter when the same hand was repeated even when different grasps were required (i.e., in condition HR). In contrast, no RT advantage was observed when participants repeated the same grasp with the opposite hand (GR). The repetition effect was accompanied by reduced fMRI activity in bilateral posterior parietal cortex (PPC), specifically in the intraparietal and superior parietal cortices. A reduction of neural activity is frequently observed when stimulus or task features are repeated and are referred to as repetition suppression effects (20–22). The authors concluded that action plans are computed independently for each hand and that, thus, PPC processes action plans separately for each effector.

These findings potentially support effector-based accounts of PPC organization which have suggested that the PPC is subdivided into lower-dimensional, effector-specific subspaces that encode motor goals in relation to a specific effector (e.g., 23, 24). However, given the primarily contralateral organization of PPC (18, 25–27), differential fMRI activation for the left and right hands might reflect effects related to body and brain symmetry rather than effector specificity (28). Indeed, fMRI activity patterns were strikingly similar in many parts of PPC when participants prepared to point to visual targets with the hand or foot of one body side (29, 30), indicative of effector-independent action representations (see also 31). In these studies, effector-specific hand and foot activity was evident only in anterior parts of PPC. Moreover, repetition suppression was evident across hand and foot in caudal parts of PPC, suggesting shared neural resources for the two limbs in the context of action planning (20). In sum, these findings suggested that PPC comprises both effector-specific and effector-independent action processing and have led to the proposal that PPC may better be viewed as organized according to functional than anatomical criteria (28). In this framework, repetition effects – both behavioral and neural (18, 19) cannot be conclusively attributed strictly to effector-specificity, but may instead at least partly reflect activation of effector-independent regions within the PPC of one hemisphere due to executing consecutive actions with limbs of one body side.

To test this alternative interpretation, we modified the paradigm of Valyear and Frey (19) to include action sequences in which the second of two consecutive actions involved repeating movements with a different effector of the same body side. Participants executed actions with hands and feet rather than just the hands, as had been done in previous studies. Following previous work, we will term the first action “prime” and the second “probe” from here on.

The movements in our tasks were hand and foot movements of both body sides in one of two directions: towards the body midline (inward) and away from the body midline (outward). Each action was cued by means of different shapes that were visually displayed on a computer screen (see Figure 1A; 19). The probe comprised the same or different effector type (hand vs. foot), the same or different body side (left vs. right), and the same or different movement direction (inward vs. outward). Thus, our task comprised a 2 × 2 × 2 factorial design that allowed dissociating effector-specific from effector-independent, hemispheric-specific repetition effects (Figure 2): Effector Type (repeat, no repeat) x Body Side (repeat, no repeat) x Movement Direction (repeat, no repeat).

**Figure 1.**
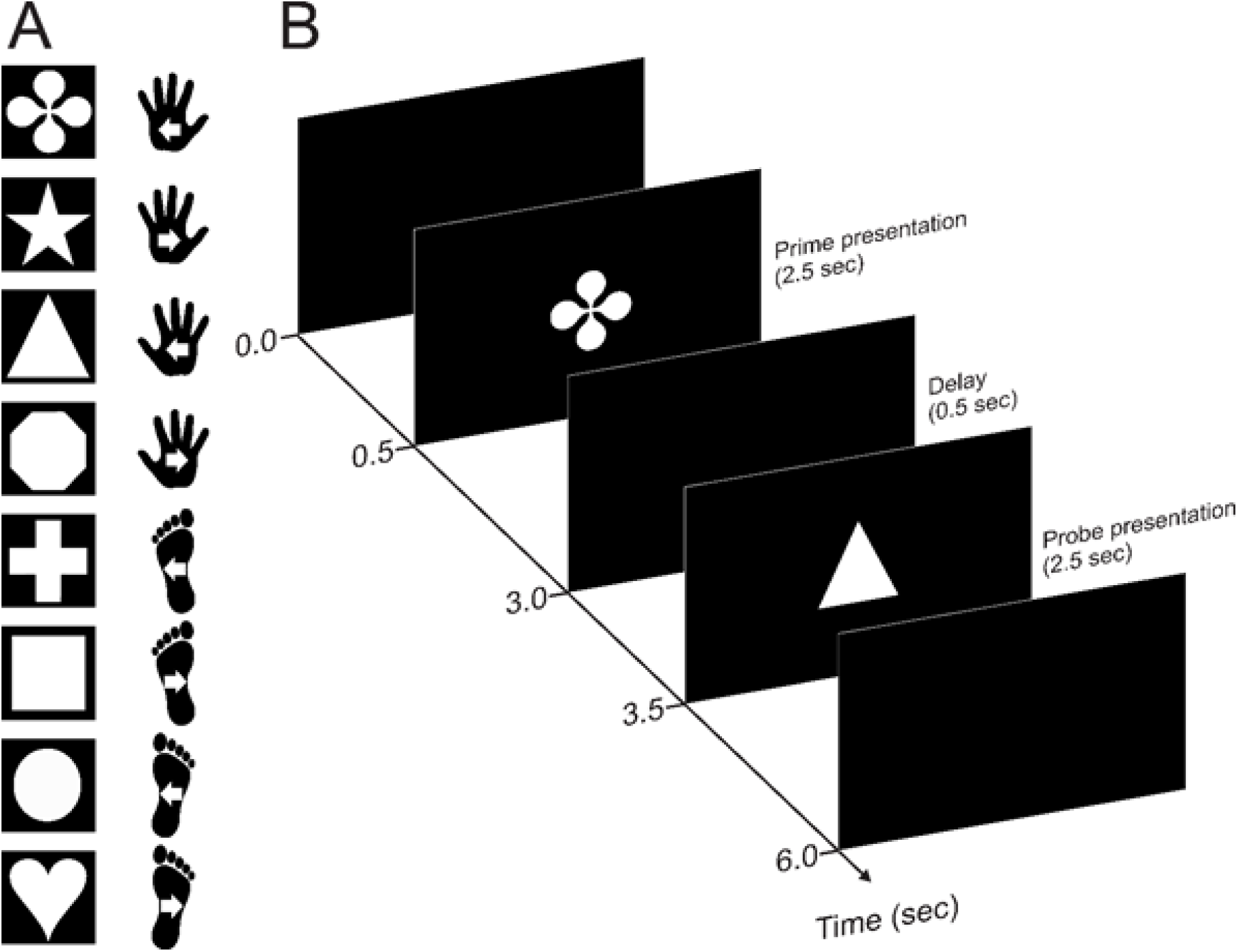
Experimental task and procedure. Participants performed two successive movements (a prime and a probe) separated by a 0.5 sec delay interval. Movements were performed in one of two directions (inward, outward) with either (i.e., left, right) hand or foot. **A** Example shape-action assignment. Actions were defined through arbitrary rules by means of different shapes visually displayed on a computer screen. Each shape corresponded to one specific action and shape-action associations were randomized across participants. Movement direction was instructed in an egocentric reference frame (i.e., in- vs. outward movement) in Experiment 1 and in an allocentric reference frame (i.e., left- vs. rightward movement) in Experiment 2. White arrows indicate movement direction. **B** Example trial sequence with event timing.

**Figure 2.**
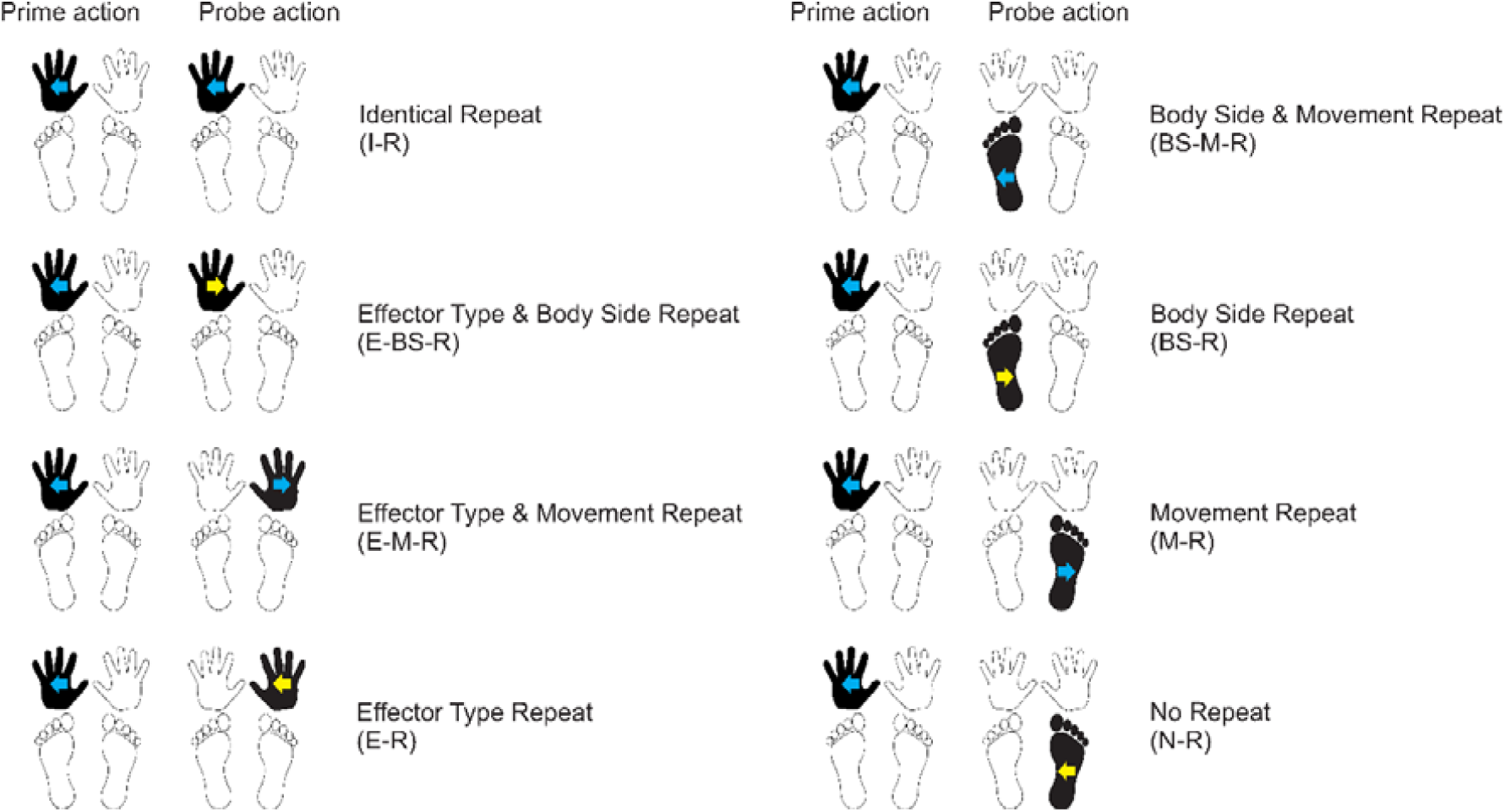
Experimental conditions of Experiment 1 (egocentric instruction). The task comprised a 2 Effector Type (Repeat, No repeat) x 2 Body Side (Repeat, No Repeat) x 2 Movement Direction (Repeat, No Repeat) design that yielded eight conditions. The left and right columns illustrate conditions in which the same (e.g., hand – hand) or different effector type (e.g., hand – foot) was used for prime and probe action, respectively. The upper and lower two rows show conditions in which the same (e.g., left – left) or different body side (e.g., left – right) was used, respectively. Conditions with same-colored (blue – blue) and different-colored arrows (blue – yellow) indicate conditions in which movements were performed in the same (e.g., outward – outward) or different (e.g., outward – inward) direction. Conditions are illustrated for a left hand outward movement as prime action but apply analogously for all other actions.

When effector type, body side, and movement direction were all repeated (identical repeat, IR), the visual cue symbol of prime and probe was also identical. For all other comparisons, prime and probe involved different symbols, because they involved a change in at least one factor and, thus, were cued by an individual shape. The IR visual cue repetition makes it difficult to interpret comparisons with the IR condition, because we cannot exclude that effects stem from visual processing of the cue and/or associative rule retrieval. In fact, the IR condition has yielded by far the fastest RT in previous studies (19). The visual confound, therefore, impedes interpretation of main effects and interactions of our factorial design. For this reason, we used as baseline the condition in which neither effector type, body side, nor movement direction were repeated (No repeat, NR); such prime-probe sequences always comprised different symbol cues. To assess repetition effects, we compared RT in the NR condition with all other conditions in dedicated contrasts.

If previously reported behavioral repetition effects reflected effector-specific representations, then our adapted design should evoke faster RT only for probe events that involved repeating the same effector type on the same body side (repeat effector type & repeat body side). Conversely, if previous repetition results were actually due to activation of effector-independent representations within one hemisphere, and not to reusing the same effector, RT should also be faster for probe events that involved repeating the same body side without repeating the same effector type (body side & movement repeat; body side repeat). Finally, our experimental design allows testing whether repeating only movement direction (movement repeat) or effector type (hand vs. foot) affects action planning (effector type & movement repeat; effector type repeat). Previous studies that involved only the hands were necessarily agnostic about this question, as it requires testing movements with different effector types, here, hand and foot.

## Experiment 1: Egocentric task instruction

### Methods

#### Participants

A previous study that involved only the two hands reported a significant repetition effect with N = 16 (19). Because we suspected that repetition effects may be smaller when instructed movements implicate the same body side but not the same effector, we defined a larger target sample size of 20 participants.

We collected data from 20 participants (1 left hander; 15 female, 5 male; mean age = 23.40 years, SD = 6.39 years, range = 19 – 49 years; mean handedness score right handers = 100.00, handedness score left hander = −50; 32). All participants were physically and neurologically healthy and reported normal or corrected-to-normal vision. They received course credit for their participation. The Bielefeld Ethics committee approved all experiments (Ethical Application Ref: 2017-114) and all participants provided written informed consent prior to participation.

#### Apparatus and stimuli

Participants sat on a chair in front of a 24” computer monitor (Dell UltraSharp U 2412M; 1920×1200 pixels; 60Hz refresh rate; height adjusted to eye level) at a distance of about 75 cm. Four square PVC blocks (width × length × height: 10 × 10 × 3 cm) with centrally embedded round pushbuttons of 7 cm diameter served as start locations for the hands and feet. The buttons were depressed by default when the limbs rested on them purely by the limbs’ weight. They were positioned such that the participants could comfortably place their hands and feet on them. The foot buttons were placed on the floor about 40 cm in front of the monitor and about 40 cm apart (center-to-center distance). The hand buttons were positioned 80 cm directly above the foot buttons on a small board. Thus, hand and foot buttons were positioned with identical distances with respect to the body and the vertical body midline.

Visual shape cues instructed the movements participants were to execute (see Figure 1A). They were colored white and sized 413 × 413 pixels. They were presented centrally on a black background (Figure 1A). Each shape defined one of eight response actions, as defined by the 3-factor design of the experiment: an inward or outward movement with respect to the body midline of about 20 cm performed with the left or right hand or with the left or right foot. On average, participants moved their limbs 16.3 cm (SD = 3.7 cm) and 16.5 cm (SD = 3.9 cm) for prime and probe actions, respectively. Movement distances were also similar across limbs, movement direction and movement phases (see Supplemental Table S1; https://osf.io/v8m9j/). Stimulus-response assignment was randomized across participants. Stimulus presentation and response recording was controlled with Matlab (The MathWorks Version R2015; Natick, MA, USA) using the Psychophysics Toolbox (33). An optical motion capture system (Visualeyez II VZ4000v, Phoenix Technologies Inc, Vancouver, BC, Canada) recorded hand and foot kinematics at 250 Hz sampling rate. We placed infrared markers on the dorsal side of each hand and foot (distal end of the third metacarpal and third metatarsal, respectively).

#### Procedure

The trial sequence is illustrated in Figure 1B. At the start of each trial, participants depressed the four response buttons with their hands and feet. After an interval of 500 ms, the prime stimulus (one of the eight shapes) appeared on the screen for 2500 ms. Participants then performed the action instructed by the stimulus. After a fixed interval of 500 ms, the probe stimulus (again one of the eight shapes) appeared on the screen for 2500 ms and participants then performed the respective probe action. After an inter-trial interval of 2000 ms, participants could initiate the next trial by depressing the four buttons. Participants received written feedback on the screen if they did not complete the respective action within the 2500 ms stimulus display time (“Too slow”) and if they released the wrong button, that is, a button of an effector that did not belong to the displayed cue (“Wrong effector”). Participants were instructed to perform inward or outward movements with the respective effector at a comfortable speed. All 64 possible prime-probe combinations were presented once within a block in a pseudorandomized order, and participants completed 10 blocks for a total of 640 trials. Due to a technical error with motion data recording, four participants performed 11 rather than 12 blocks.

Participants performed 2-3 practice blocks prior to the experimental blocks until they reported to be sufficiently familiar with the stimulus-response associations. The trial structure was identical to the experimental trials except that only one stimulus was shown and participants performed only one action per trial. If participants used the wrong effector, an error message indicating the correct response was displayed. Each stimulus was repeated 8 times within a block, yielding 64 trials per block. The entire experiment took about 3 hours to complete.

#### Data processing and analysis

We processed kinematic data using custom-written Matlab scripts. We interpolated missing data points using the spring metaphor method in the inpaint_nans function (34) and smoothed the data using a second-order Butterworth filter with a cut-off frequency of 6 Hz. We determined movement onset as the time point at which the vectorial velocity of a marker exceeded 50 mm/s and response time (RT) as the time between probe stimulus onset and movement onset. We excluded trials in which no data was recorded and trials in which trajectories could not be reconstructed due to too many missing data points (2.3%, min. 0.2%, max. 17.6% per participant). We removed trials in which participants performed the wrong prime or probe action (8.8%, min. 2.2%, max. 18%) and trials in which RT was faster than 200ms or slower than 1500ms (1.2%, min. 0%, max. 5.3%). We also considered analyzing repetition effects on errors in action choice. However, trials in which participants committed an error during the probe phase were very rare (Wrong effector: 1.1%, Wrong Body Side: 1.3%, Wrong movement direction: 0%), and hence conclusions about this aspect of our data are problematic.

#### Statistical approach

We analyzed our data in R (35) using Bayesian regression models created in Stan (http://mc-stan.org/) and accessed with the package brms version 2.13.0 (36).

In a first step, we fitted Bayesian regression models with RT as dependent variable and the within-subject variables Effector (hand, foot), Body Side (left, right), and Movement Direction (inward, outward), separately for prime and probe actions, to assess whether RT differed for the different movement types and, thus, confounded cross-limb comparison.

For our main analysis, we fitted a model for the dependent variable RT only for probe actions that included the within-subject categorical variables Effector Type Repeat (repeat, no repeat), Body Side Repeat (repeat, no repeat), and Movement Direction Repeat (repeat, no repeat) as independent predictors. We set orthogonal contrasts using the set_sum_contrasts() command in afex version 0.28-0 (37). The model was specified as RT ∼ Effector Type Repeat * Body Side Repeat * Movement Direction Repeat + (1 | participant), thus capturing all main effects and interactions of our design. For all models, we included only random intercepts, because models did not reliably converge when random slopes were included. We used a shifted log-normal distribution to estimate RTs. To improve convergence and guard against overfitting, we specified mildly informative, conservative priors for population-level (i.e., fixed) effects. The estimation of parameters’ posterior distributions were obtained by Hamiltonian Monte-Carlo sampling with 4 chains, 1,000 sample warmup, and 11,000 iterations, and checked visually for convergence (high ESS and Rhat ≈ 1). To analyze and describe the posterior distributions, we report the median as a point-estimate of centrality and the 95% credible interval (CI) computed based on the highest-density interval (HDI) to characterize the uncertainty related to the estimation. As an index of the credibility of a given effect, we tested whether the HDI excluded a region of practical equivalence (ROPE) of +/-0.1 effect sizes around 0 (38, 39). If the HDI is completely outside the ROPE, the null hypothesis for this effect is rejected. If the ROPE completely covers the HDI (i.e., all most credible values of a parameter are inside the ROPE), the null hypothesis is accepted. This approach is conservative because it requires not only that the HDI exclude 0, but in addition that no overlap exists between the HDI and the ROPE interval.

#### Results

RT was strikingly similar across the different movement types (i.e., hand and foot; left and right body side, inward and outward movement direction) for both prime and probe actions (Supplemental Table S2; https://osf.io/g98er/). Bayesian analysis confirmed that the HDI was inside the ROPE for all predictors for both prime and probe actions. This result demonstrates that none of the parameters affected RT (Supplemental Figure S3; https://osf.io/hy5zb/). Together, these results thus indicate that there are no baseline RT differences that might influence our main analysis.

RT for probe actions was fastest when effector type, body side, and movement direction were all repeated (identical repeat, IR; median = 525 ms, 95% HDI [503, 548]) and slowest when none of the three tested criteria was repeated (no repeat, NR; median = 790 ms, 95% HDI [750, 831]). Compared with NR, RT was faster when effector type and body side were repeated (median = 716 ms, 95% HDI [681, 752]); recall that this factor combination reflects true effector-specificity, because the combination of repeated effector type and body side implies re-use of a given effector. In contrast, RT was similar when only body side but not effector was repeated (body side repeat & movement direction repeat: median = 781 ms, 95% HDI [741, 822]; body side repeat: median = 779 ms, 95% HDI [739, 819]). Importantly, this pattern was similar for hand and foot movements (Figure 3). When we extended the Bayesian regression model by a factor Effector Type (hand, foot; complete model: RT ∼ Effector Type Repeat * Body Side Repeat * Movement Direction Repeat * Effector Type), the HDI for Effector Type and all its interactions were inside the ROPE. Thus, Effector Type did not statistically modulate the repetition effect (Supplemental Figure S4; https://osf.io/yt4xk/).

**Figure 3.**
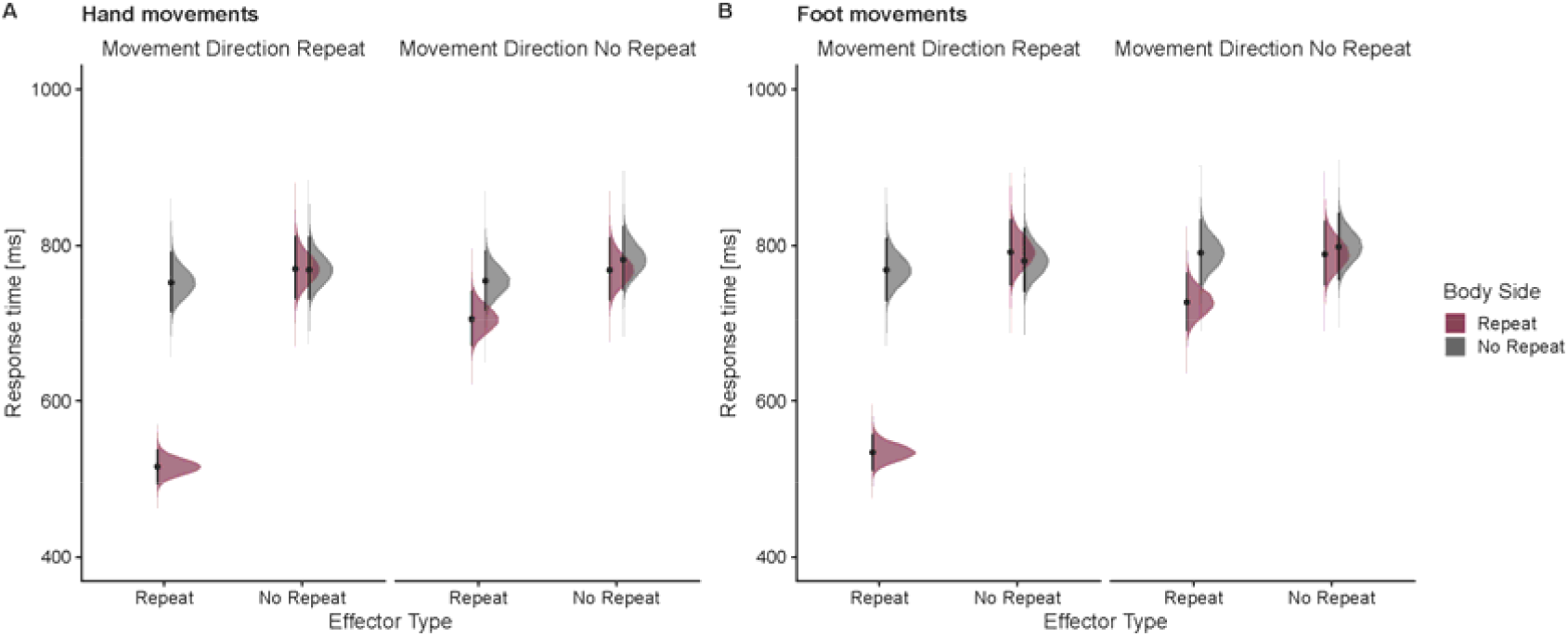
Response times (ms) for probe actions as a function of effector type, body side, and movement repeat, separately for hand movements (A) and foot movements (B) in Experiment 1. Areas show the posterior distributions of the estimated marginal means, black dots represent the medians and error bars the 95% highest density interval (HDI).

To distinguish between effector- and effector-independent hemispheric-specific repetition effects, we calculated contrasts between the posterior distributions of NR and conditions that involved repetition of the same effector type and/ or body side (i.e., IR, EBSR, BSMR, BSR; Figure 4 upper four rows). Repeating the action (IR) resulted in a substantial decrease of RT compared to NR (NR – IR: median difference = 265 ms, 95% HDI [244, 286]; 0% of HDI in ROPE). However, as mentioned previously, these differences cannot be separated from visual processing and associative rule retrieval. RT was also faster than NR when effector and body side were repeated (NR – EBSR, median difference = 74 ms, 95% HDI [60, 89]; 0% of HDI in ROPE). In contrast, comparisons of conditions that involved repetition of the same body side but not effector type did not yield any credible RT differences against NR (NR – BSMR: median difference = 9 ms, 95% HDI [−5, 24]; NR – BSR: median difference = 11 ms, 95% HDI [−3, 26]; Figure 4, rows 3 and 4) and 88% and 81% of the HDIs overlapped with the ROPE, respectively. In addition, comparison of conditions that did vs. did not include an effector-specific repeat were credibly different, suggesting a critical influence of repeating a specific effector rather than just any limb of the given body side (BSMR – EBSR: median difference = 65 ms, 95% HDI [51, 79]; 0% of HDI in ROPE; BSR – EBSR: median difference = 63 ms, 95% HDI [49, 77]; 0% of HDI in ROPE; Figure 4 bottom two rows). These findings lend support to the view that repetition effects in effector choice reflect effector-specific rather than effector-independent hemisphere-specific coding of action representation. Lastly, we did not find evidence for RT improvement when only the effector type was repeated (NR – EMR: median difference = 30 ms, 95% HDI [15, 44]; 17% of HDI in ROPE; NR – ER: median difference = 18 ms, 95% HDI [4, 33]; 57% of HDI in ROPE; Figure 4, rows 5 and 6) or when only movement direction was repeated (NR – MR: median difference = 16 ms, 95% HDI [2, 31]; 64% of HDI in ROPE; Figure 4 row seven).

**Figure 4.**
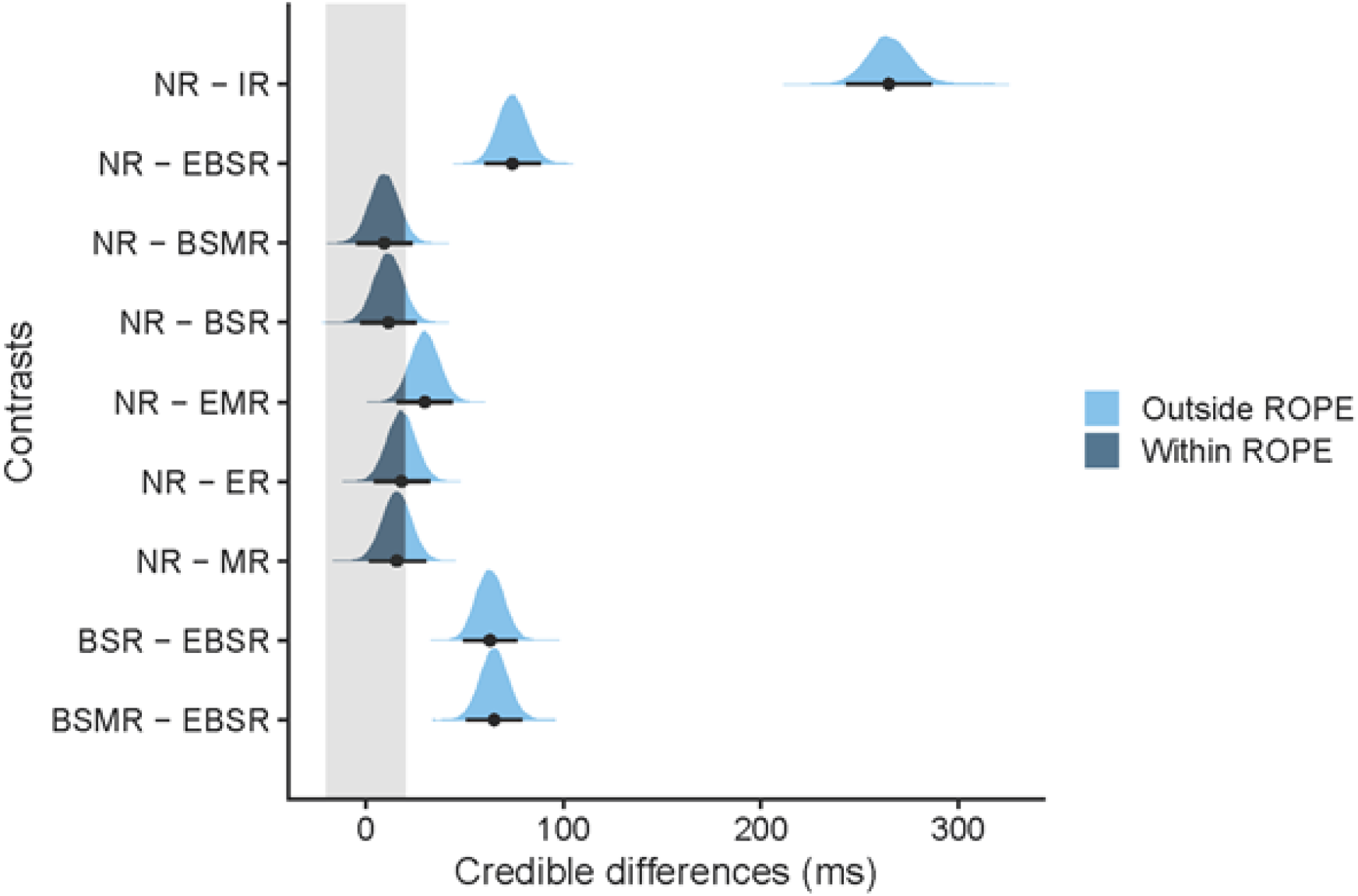
Contrasts of posterior distributions in Experiment 1. Black dots represent the medians and error bars the 95% highest density interval (HDI). The grey shaded area indicates the region of practical equivalence (ROPE) of +/-0.1 effect sizes around 0 [−20, 20]. Portions of the distributions outside and inside the ROPE are shown in light blue and dark blue, respectively. NR = No repeat, IR = identical repeat, EBSR = effector type repeat and body side repeat, BSMR = body side repeat and movement direction repeat, BSR = body side repeat, EMR = effector type repeat and movement direction repeat, ER = effector type repeat, MR = movement direction repeat.

By reviewer request, we re-analyzed our experiment with frequentist statistics using repeated measures ANOVA. This analysis rendered qualitatively similar results (see Supplemental Table S5; https://osf.io/78ubr/).

## Experiment 2: Allocentric task instruction

In Experiment 1, repeating using the same effector repeatedly resulted in a substantial RT benefit even when the performed actions were different (e.g., left hand inward movement – left hand outward movement). In contrast, movements of two different effectors of one body side (e.g., left foot – left hand) did not yield a credible decrease in RT. These findings suggest that repetition effects in limb selection reflect effector-specific rather than effector-independent hemispheric-specific action representations.

Movements of homologous limb pairs, such as the two hands, are more stable and accurate when they are symmetrical with respect to the sagittal body midline than when the two limbs have to coordinate non-symmetrically. This symmetry advantage is known as the “egocentric constraint”. In contrast, movements of non-homologous limb pairs (e.g., one hand and one foot are more stable and accurate when the two limbs move in the same direction in extrinsic space, known as the “allocentric constraint” (cf. 40). In Experiment 1, participants were instructed to perform inward and outward movements, that is, instructions were formulated egocentrically. This choice of instructions may have penalized movements performed with non-homologous and, hence, prevented the expression of body side-specific repetition effects. To test this conjecture, we instructed participants allocentrically in Experiment 2: they were now asked to perform left- and rightward-directed movements in response to the visual stimuli.

### Methods

#### Participants

20 individuals from Bielefeld University (1 left hander; 8 female, 12 male; mean age = 22.90 years, SD = 1.94 years, range = 20 – 28 years; mean handedness score right handers = 97.53, SD = 7.63, handedness score left hander = −43; 32) participated in Experiment 2. None of them had participated in Experiment 1. All participants were physically and neurologically healthy, reported normal or corrected-to-normal vision, and received course credit in exchange for their participation.

#### Apparatus, stimuli, and procedure

The apparatus, stimuli, and procedure were identical to Experiment 1 except that movements were instructed and trained as leftward and rightward (allocentric response coding), as opposed to inward or outward (egocentric response coding). On average, participants moved their limbs 18.5 cm (SD = 2.8 cm) and 18.9 cm (SD = 3.8 cm) for prime and probe actions, respectively. Movement distances were also similar across limbs, movement direction, and movement phases (see Supplemental Table S6; https://osf.io/8db9m/).

#### Data processing and analyses

Data processing and analysis were analogous to Experiment 1. We excluded trials in which no data was recorded and trials in which trajectories could not be reconstructed due to too many missing data points (1.7%, min. 0.3%, max. 6.1% per participant). We removed trials in which participants performed the wrong prime or probe action (9%, min. 1.3%, max. 25.7% per participant) and trials in which RT was faster than 200ms or slower than 1500ms (1.6%, min. 0%, max. 8% per participant). Trials in which participants committed an error during the probe phase were similarly rare as in Experiment 1 (Wrong effector: 1.0%, Wrong Body Side: 1.7%, Wrong movement direction: 1.0%), hence we also omitted analyzing repetition effects on action choice in Experiment 2.

#### Results

RT was similar across body side (left, right) and movement direction (leftward, outward). In contrast to Experiment 1, RT was faster for hand than foot movements for both prime and probe actions (Supplemental Table S7; https://osf.io/p9hvy/). Bayesian analysis showed that only the HDI for the predictor Effector Type was outside the ROPE for both prime and probe actions (prime: median = 39 ms, 95% HDI [34, 45]; probe: median = 40 ms, 95% HDI [35, 45]), whereas the HDI for the other predictors were completely or almost completely inside the ROPE (Supplemental Figure S8; https://osf.io/69sru/). Thus, the RT difference between hands and feet resembled an offset that did not modulate any other effects present in our data.

With respect to repetition effects, results were remarkably similar to those of Experiment 1. RT for probe actions was fastest when effector type, body side, and movement direction were all repeated (identical repeat, IR; median = 525 ms, 95% HDI [497, 553]). Compared to NR (median = 757 ms, 95% HDI [708, 809]), RT as again faster when effector type and body side were repeated (median = 697 ms, 95% HDI [654, 743]). In contrast, RT was relatively similar when only body side but not effector was repeated (body side repeat & movement direction repeat: median = 754 ms, 95% HDI [705, 805]; body side repeat: median = 754 ms, 95% HDI [704, 804]). Again, this pattern was similar for hand and foot movements (Figure 5). A Bayesian regression model extended with factor Effector Type (hand, foot; complete model: RT ∼ Effector Type Repeat * Body Side Repeat * Movement Direction Repeat * Effector Type) confirmed that the HDI for the factor Effector Type was outside the ROPE (median = 41 ms, 95% HDI [36, 46]). Importantly however, the HDI for all interactions with the factor Effector Type were completely inside the ROPE, indicating that this factor did not modulate the repetition effect (Supplemental Figure S9; https://osf.io/6vrxa/).

**Figure 5.**
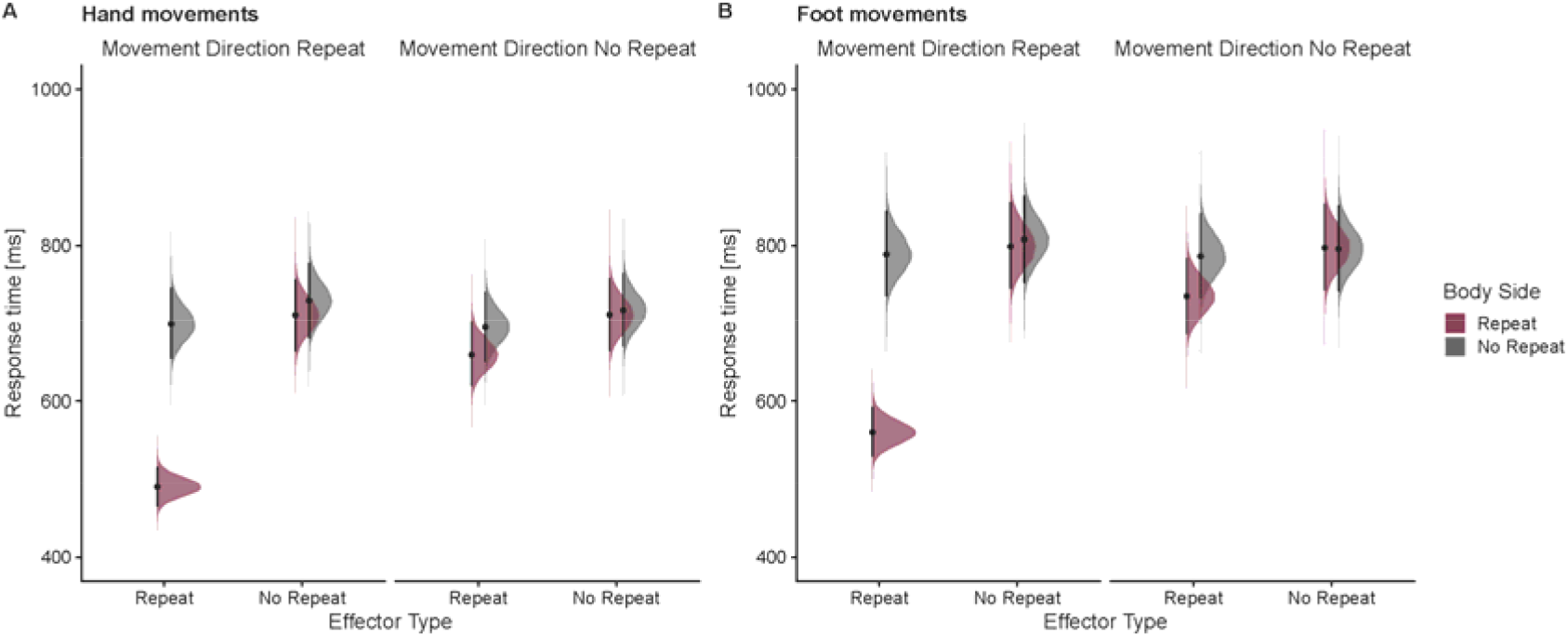
Response times (ms) for probe actions as a function of effector type, body side, and movement repeat, separately for hand movements (A) and foot movements (B) in Experiment 2. Areas show the posterior distributions of the estimated marginal means, black dots represent the medians and error bars the 95% highest density interval (HDI).

Again, we calculated contrasts between the posterior distributions of NR and conditions that involved repetition of the same effector type and/ or body side to distinguish between effector- and effector-independent hemispheric-specific repetition effects (i.e., IR, EBSR, BSMR, BSR; Figure 6 upper four rows). RT was substantially faster in IR (median difference = 231 ms, 95% HDI [207, 257]; 0% of HDI in ROPE), but as in Experiment 1, this contrast may be strongly affected by factors that are not of interest in our study. RT was also faster when a specific effector was repeated (EBSR, median difference = 60 ms, 95% HDI [46, 73]; 0% of HDI in ROPE). In contrast, there were no credible RT differences for repetition of body side (NR – BSMR: median difference = 3 ms, 95% HDI [−10, 16]; NR – BSR: median difference = 3 ms, 95% HDI [−11, 16]), and the 95% HDI of the respective credible difference distributions were fully inside the ROPE. In addition, comparison of specific effector repetition (i.e., repeating effector and body side) and repeating only body side confirmed credible differences between these conditions (BSMR – EBSR: median difference = 57 ms, 95% HDI [43, 71]; 0% of HDI in ROPE; BSR – EBSR: median difference = 57 ms, 95% HDI [43, 71]; 0% of HDI in ROPE; Figure 6, bottom two rows). Finally, there were no credible effects for repeating only effector type but not body side (NR – EMR: median difference = 14 ms, 95% HDI [0, 27]; 72% of HDI in ROPE; NR – ER: median difference = 17 ms, 95% HDI [4, 31]; 59% of HDI in ROPE; Figure 6 rows five and six) and when repeating only movement direction (NR – MR: median difference = −12 ms, 95% HDI [−26, 2]; 78% of HDI in ROPE; Figure 6 row seven). In sum, these results confirm the findings of Experiment 1, and the two experiments together reinforce the view that repetition effects in limb selection originate from effector-specific processes.

**Figure 6.**
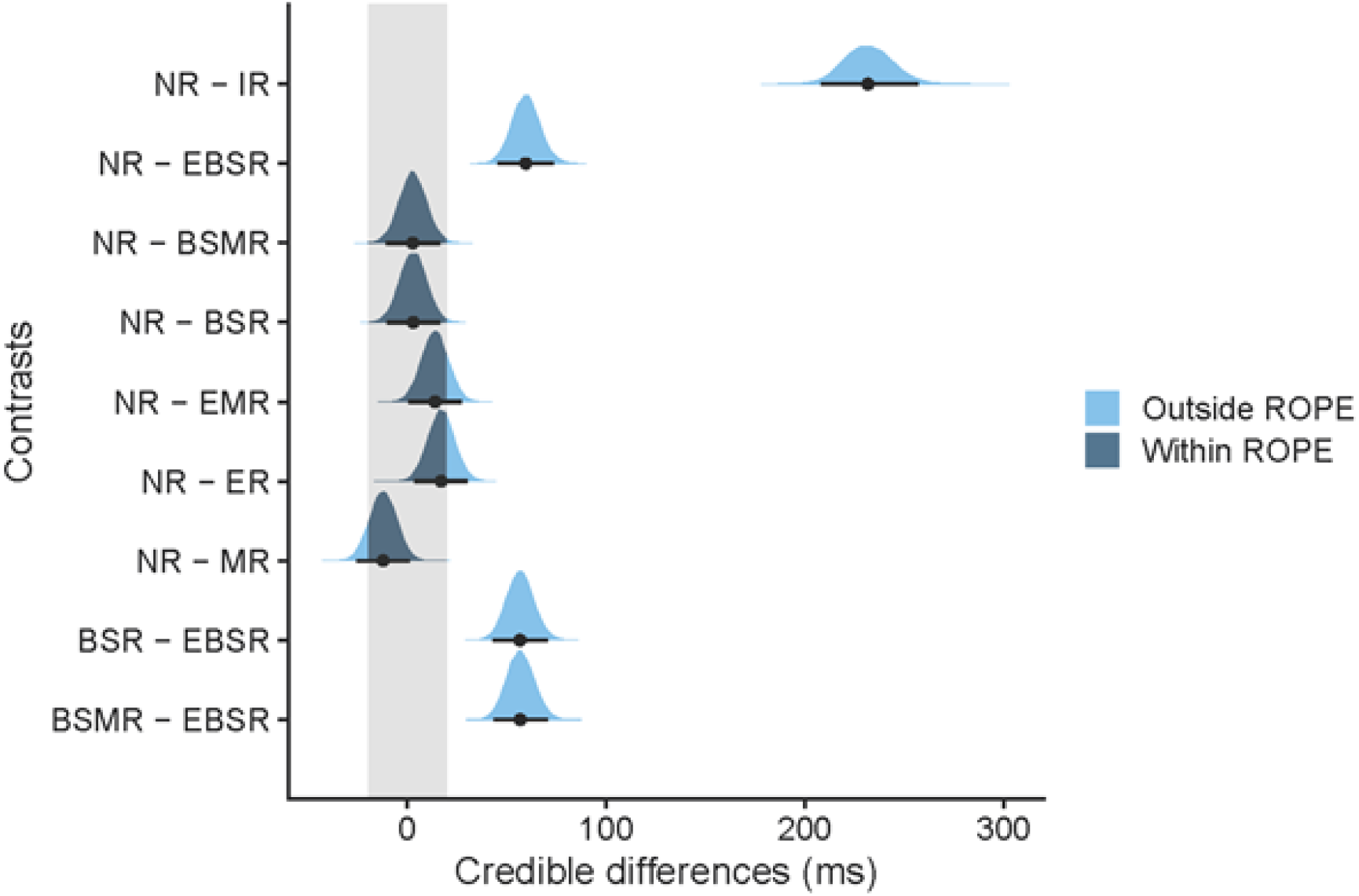
Contrasts of posterior distributions in Experiment 2. Black dots represent the medians and error bars the 95% highest density interval (HDI). The grey shaded area indicates the region of practical equivalence (ROPE) of +/-0.1 effect sizes around 0 [−20, 20]. Portions of the distributions outside and inside the ROPE are shown in light blue and dark blue, respectively. NR = No repeat, IR = identical repeat, EBSR = effector type repeat and body side repeat, BSMR = body side repeat and movement direction repeat, BSR = body side repeat, EMR = effector type repeat and movement direction repeat, ER = effector type repeat, MR = movement direction repeat.

As for Experiment 1, we re-analyzed our data with repeated measures ANOVA. As before, the results were qualitatively similar (see Supplemental Table S10; https://osf.io/z632x/).

#### Modeling

We employed computational modeling to examine the source of the (effector-specific) repetition effects. We analyzed reaction times and accuracy data of both experiments (n = 40) within the framework of drift-diffusion models (DDM; 41). According to DDM, decision making in binary decision tasks can be regarded as an accumulation-to-threshold mechanism in which activity related to the different response options builds up as a function of the evidence for or against those choices until one of two opposing decision bounds is reached. These models can explain both accuracy patterns and response latencies across a variety of different tasks (41–46).

Moreover, neural activity in the posterior parietal cortex seems to reflect the accumulated evidence in perceptual decision-making tasks (44). One explanation is that repetition effects are a consequence of persistent changes in baseline activity within neurons encoding effector-specific action plans. Accordingly, sustained elevated neural activity for recently selected effectors would result in faster action initiation (13, 18). Within the DDM framework, this can be modeled as a shift of the starting point of the evidence accumulation process towards the repeated action in repeat trials as compared to non-repeat trials (starting point model). The shift reduces the distance from the starting point to one of the decision bounds in effector-repeat trials, so that less evidence accumulation is required until the bound is reached and, accordingly, less time will pass until the associated movement is initiated.

Repetition effects may also emerge through a higher drift rate in repeat over non-repeat trials (drift rate model). The drift rate is a measure of the speed of information processing and higher values indicate a faster rate of evidence accumulation such that the bound is reached faster. Changes in either parameter can explain sequential effects (cf. 47).

In addition to these two model variants, we fitted a model in which both starting point and drift rate varied per repeat and non-repeat condition (full model). Finally, we considered a model in which starting point and drift rate were fixed across all conditions to assess whether any of the two parameters of interest is necessary to explain behavior (baseline model).

We fitted each of the four model variants with the maximum-likelihood approach implemented in the software fast-dm (48, 49). All model variants included the parameters drift rate *v*, starting point *z*_*r*_, accumulation threshold *a*, and non-decision time *t*_*0*_. The four model variants differed in terms of which parameters were allowed to vary across Effector Type, Body Side, and Movement Direction conditions: In the full model, drift rate and starting point were allowed to vary simultaneously. In the drift rate model, only the drift rate, and in the starting point model, only the starting point could vary between conditions. In the baseline model, no parameter was allowed to differ between conditions. In all model variants, inter-trial variabilities of drift rate and starting point were fixed at zero, because simulations show weak recovery for these parameters, especially in smaller data sets (50). However, we included variability of the starting point *s*_*t0*_, as this can counteract data outliers (51). We fitted each model to individual participants’ probe stimuli RT data. We only included trials that followed a correct response to the preceding prime stimulus. Otherwise, outlier criteria remained the same as in the main analysis, with the difference that for DDM analysis, both correct and incorrect responses were included; this is necessary and adequate because the DDM estimates RT distributions for both correct and incorrect responses. The models’ upper response thresholds were associated with a correct response and the lower threshold with any possible incorrect response option.

We compared the BIC/AIC weights of the individual model fits by converting the Bayesian information criterion (BIC) and Akaike information criterion (AIC) for each model (52) to determine the best model in terms of goodness of fit and parameter parsimony. The starting point model provided the best fit, followed by the drift rate and full model, while the baseline model provided the worst fit (Table 1). Although BIC and AIC ordered drift rate and full models differently, the two criteria were consistent concerning the starting point and baseline model.

**Table 1.**
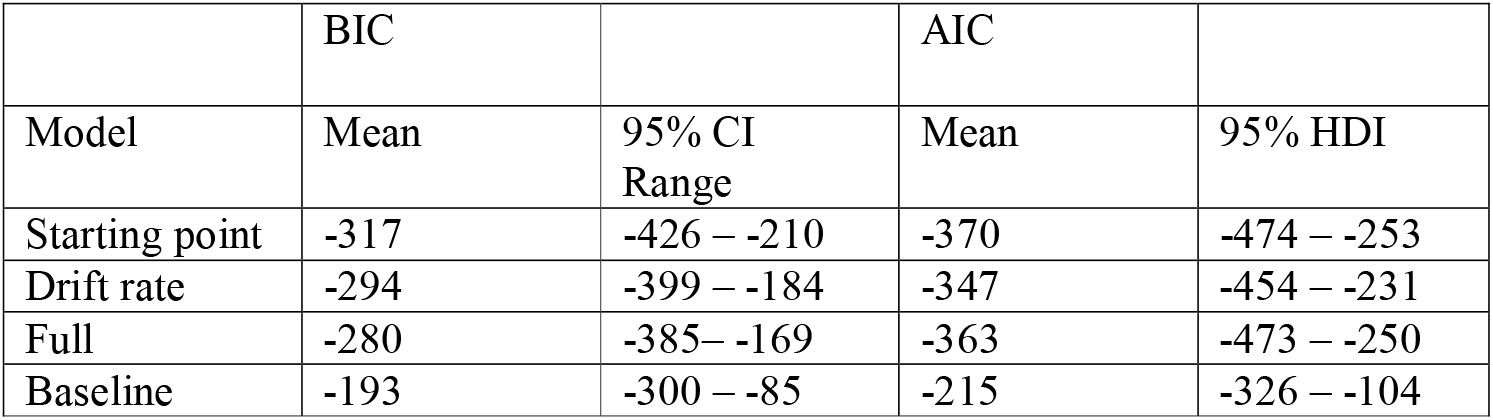
BIC and AIC calculated for different DDM. Estimated mean values and 95% HDI using Bayesian regression (BIC/ AIC ∼ model variant + (1|participant)).

|  | BIC |  | AIC |  |
| --- | --- | --- | --- | --- |
| Model | Mean | 95% CI Range | Mean | 95% HDI |
| Starting point | -317 | -426 – -210 | -370 | -474 – -253 |
| Drift rate | -294 | -399 – -184 | -347 | -454 – -231 |
| Full | -280 | -385 – -169 | -363 | -473 – -250 |
| Baseline | -193 | -300 – -85 | -215 | -326 – -104 |

We further scrutinized the relevance of the drift rate parameter by comparing the model parameters of the full model. Individual drift rates were higher for the identical repeat (IR) condition compared to all other conditions (Figure 7A). Contrasts between the posterior distributions showed credible differences between NR and IR (mean difference = −1.34, 95% HDI [−1.61, −1.06]; 0% of HDI in ROPE; Figure 7C). However, this high drift rate is likely due to facilitated processing of perceptual features, given the previously discussed visual symbol repetition (53). Importantly, drift rate was similar for the effector repeat condition (EBSR) and the NR condition (mean difference = 0.16, 95% HDI [−0.12, 0.44]; 36% of HDI in ROPE; Figure 7C), indicating that repetition effects do not reflect a change in drift rate. In contrast, starting points were shifted towards the decision bound for the IR and the effector repeat (EBSR) condition (Figure 7B). Contrasts between the posterior distributions showed credible differences between NR and IR (mean difference = −0.35, 95% HDI [−0.38, −0.32]; 0% of HDI in ROPE) and between NR and EBSR (mean difference = −0.13, 95% HDI [−0.16, −0.10]; 0% of HDI in ROPE; Figure 7D). Thus, even if we allow free fitting of drift rate for each condition, this parameter does not account for repetition effects in motor performance. These results are in line with the superior fit of the starting point model and indicate that repetition effects likely arise due to a change in baseline activity that is exclusive to a specific effector.

**Figure 7.**
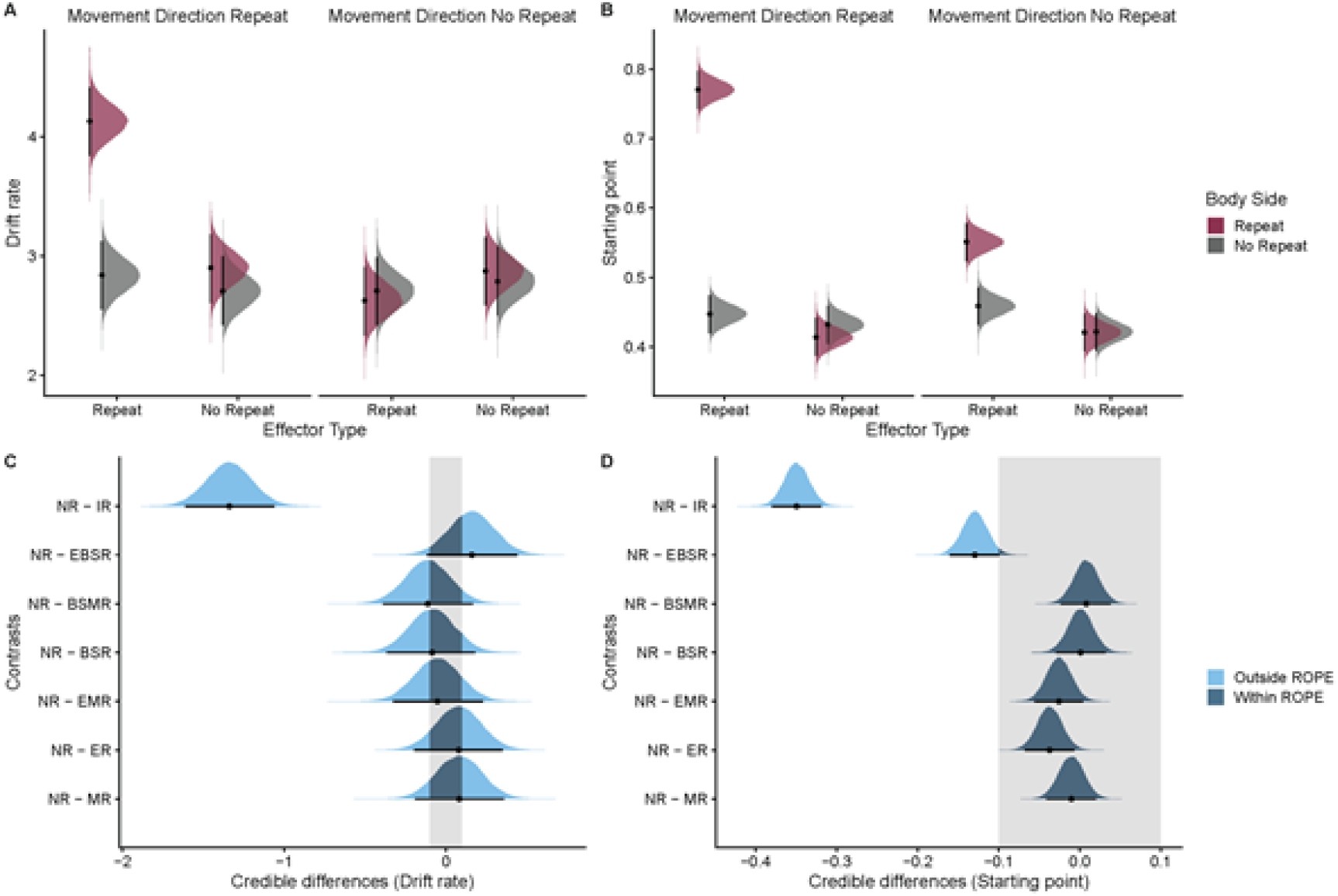
DDM parameter estimates. Posterior distributions of the estimated marginal means of drift rates (A) and starting points (B) for probe actions as a function of effector type, body side, and movement repeat calculated from the DDM (full model). (C,D) Contrasts of posterior distributions between the no repeat and the other conditions for drift rates (C) and starting points (D). Black dots represent the means and error bars the 95% highest density interval (HDI).The grey shaded area indicates the region of practical equivalence (ROPE) of +/-0.1 effect sizes around 0. Portions of the distributions outside and inside the ROPE are shown in light blue and dark blue, respectively.

## Discussion

The aim of the present study was to test whether repetition effects in limb selection reflect effector- or effector-independent hemisphere-specific action representations. Participants performed movements with hands and feet in one of two directions. Successive actions were initiated faster when the same effector was moved repeatedly (e.g., left hand – left hand), even when those actions involved different movement directions. Conversely, moving two effectors of the same body side (e.g., left foot – left hand), that is, a body side repetition, did not yield any significant RT benefit. This result pattern was evident irrespective of whether movement direction was instructed egocentrically (inward vs. outward) or allocentrically (left vs. right).

These findings are consistent with the view that current limb choice is influenced by limb choice in the recent past, and that successive actions can be initiated faster when an effector is employed repeatedly (13, 19), possibly through persistent changes in baseline activity of the neural assembly responsible for the given effector, as suggested by our computational modeling approach. Importantly, our findings confirm that repetition effects arise from effector-specific processing and do not support the alternative that repetition effects are based on effector-independent processing of the same body side, which are largely controlled by one hemisphere.

Multiple studies have demonstrated that previously performed actions can bias current actions. Such motor hysteresis effects have been observed across a variety of different tasks. Besides hand choice (11, 13–15), motor hysteresis is also known to influence grasp height (54–56), grasp orientation (57–59), and reach trajectories (60, 61) such that current actions tend to be more similar to previously executed actions.

The presence of such history effects is typically explained in terms of computational gains underlying action planning. It is commonly assumed that it is computationally more efficient for the sensorimotor system to recycle motor parameters of previously specified action plans than to create them anew for every single movement (16, 17, 62). Consistent with this view, previous work has shown that actions are initiated faster when an effector is used repeatedly (13, 19), and this RT benefit was accompanied by reduced fMRI activity in bilateral posterior parietal cortex (18).

However, previous studies have consistently used homologous effectors, usually the hands, in their experiments. Thus, a non-excluded possibility so far had been that repetition effects do not originate from repeatedly using one specific effector but, instead, from an effector-independent hemisphere-specific advantage, which would, of course, also benefit action repetitions of an identical effector.

Our present results support the previously insufficiently backed-up conclusion that repetition effects in limb choice paradigms reflect processing advantages at the level of specific effectors. In turn, our results do not provide any evidence for a role of processes that encompass effector-independent coding within one hemisphere. This is because a reaction time benefit was only present when the same effector was moved twice, but not two different effectors of one body side were moved consecutively.

Our study, thus, suggests that repetition effects result from effector-specific processing, presumably in PPC (18, 23, 24, 63, 64). The concept of effector-specific PPC organization has mainly originated from neurophysiological studies in macaque monkeys that reported differential neuronal activity for hand vs. eye movements. For instance, the macaque lateral intraparietal area (LIP) is more active during saccade than reach planning, and the parietal reach region (PRR) shows the opposite activity pattern (4, 65, 66). Several fMRI studies have revealed considerable overlap for saccade and reach planning in human PPC (67, 68), and these results have challenged strict effector-related accounts of PPC. Still, effector-specific activity for saccades vs. reaches could be decoded with multivoxel pattern analysis in several human PPC regions such as the superior parietal-occipital cortex (SPOC) and the posterior intraparietal sulcus (69).

Nevertheless, the reliance of effector-specificity accounts on the dissociation of hand and eye movements has been challenged (cf. 28), based on the argument that eye and hand movement serve different, namely perceptual-exploratory vs. manipulatory, purposes. Studies that contrasted planning-related activity of eye, hand, and foot movements have produced a more differentiated picture of PPC organization (20, 29, 30). Specifically, these studies have revealed a caudo-rostral gradient for eye vs. limb movements and a lateral-to-medial gradient for hand vs. eye vs. foot movements in anterior PPC, combined with mostly common activation patterns and voxel patterns for multiple effectors effectors in posterior, and differentiated activity in anterior PPC. Moreover, some anterior PPC regions appeared to code one effector against the others. Based on these dichotomous representations, the authors proposed that these anterior regions probably play a role in effector selection, rather than representing one specific effector. The fact that the RS effects accompanying the RT benefit in effector repeat trials described by Valyear and Frey (18) were also partially located in regions of PPC exhibiting hand-specific activation corroborates this interpretation.

According to the Affordance Competition Hypothesis (1, 2, 70), the brain simultaneously specifies multiple actions that are presently available in the environment and decides between them through continuous competition of the neural activity related to each action within a distributed fronto-parietal network. Once the neural activity for one of the potential actions reaches a threshold, its activity is further increased while the activity of competing actions is inhibited. This perspective is in line with computational models that conceptualize decision making as an accumulation-to-threshold mechanism in which activity related to different response options builds up as a function of the evidence for each choice until a threshold is reached (41–46). Our results from computational modeling further reinforce such an interpretation. Specifically, we found that the starting point parameter was shifted towards the decision bound specifically for the condition involving effector repetition (i.e., IR, EBSR). While the shift in starting point in combination with the higher drift rate in the IR condition might largely reflect facilitated perceptual processing due to symbol repetition, it cannot readily explain the starting point shift in the EBSR condition, which does not involve symbol repetition. Rather we contend that this finding fits the idea that effector-specific RT advantages would result from changes in activity within neural populations in anterior PPC that are distinctly dedicated to a given effector. Once an effector-specific population has been activated, neural activity that remains within this particular circuit would support the building of new activity, while alternative effectors would still be suppressed by the activity left-over from the previous movement. Accordingly, evidence for the recently selected effector would be closer to the threshold, yielding shorter RTs in effector repetition trials.

In this context, it is noteworthy that other interpretations of motor repetition effects have been put forward. For example, Dixon et al. (59) showed that repetition effects in grasping were linked to object identity and persisted across intervening trials that involved different grasps, suggesting that repetition effects reflect memory retrieval processes (71). There is also evidence that hysteresis effects can transfer from one hand to the other (59, 72), indicative of an involvement of effector-independent representations in repetition effects (73). Why such cross-hand advantages arise in some, but not other studies (such as in our present one), is currently unresolved. Finally, the time course of hysteresis effects appears to depend on the task specifics and can range from below one second (60) to several seconds (16, 18, 19, 59). Thus, it is possible that the repetition effects observed in different tasks reflect different underlying processes (see also 74). It is conceivable, then, that the exclusiveness of effector-specific repetition effects identified in our present experiments may not generalize to all motor contexts.

An important aspect of our current results is the independence of the repetition effects we observed from the reference frames that specified the instructed movements in the two experiments. Although egocentric coding exhibits higher activity in parietofrontal cortex, and allocentric coding exhibits higher activity in early visual cortex, neural activity for allocentric and egocentric mechanisms for reach target coding also shows considerable overlap (75). Moreover, the repetition effects in the present study and previous work (18, 19) likely reflect effects related to effector choice and may, thus, not be directly linked to processes that involve coordinate transformations – a hypothesis that will require further exploration but has received some support from limb choice tasks in the tactile domain (76).

To summarize, we demonstrate that the successive re-use of an identical effector, but not of effectors that belong to the same body side, results in a substantial decrease in the time required for movement initiation. These findings are consistent with the view that current limb choice is influenced by choices in the recent past. They lend support to the widely held view that repetition effects in limb choice reflect effector-specific rather than effector-independent hemisphere-specific coding.

## Acknowledgments

We thank Tobias Oehler and Florian Grünendahl for help with data acquisition and Conrad Alting for help with programming of the experiments. Data and code for the present paper are available at the Open Science Framework website https://osf.io/h5zrb

## Disclosures

No conflicts of interest, financial or otherwise, are declared by the authors.

## Author contributions

C.Se. and T.H. conceived and designed research; C.Se. performed experiments; C.Se and C.Sc. analyzed data; C.Se., C.Sc., and T.H. interpreted results of experiments; C.Se. prepared figures; C.Se. drafted manuscript; C.Se., C. Sc., and T.H. edited and revised manuscript; C.Se., C.Sc., and T.H. approved final version of manuscript.

## Notes

https://osf.io/h5zrb/

### Competing Interest Statement

The authors have declared no competing interest.

